# The fate of a dynasty: Population genomics uncovers the demographic history of *Ardea insignis*, one of the rarest bird species in the world

**DOI:** 10.64898/2026.08.04.742811

**Authors:** Martin Kapun, Tshering Tobgay, Alexandra Wanka, Wolfgang Fiedler, Tricia C. Goulding, Andreas Kroh, Luise Kruckenhauser, Samten Leki, Thinley Phuntsho, Marcela Suarez-Rubio, Sonam Tshering, Swen C. Renner

**Affiliations:** Natural History Museum Vienna, Central Research Labs, Burgring 7, 1010 Vienna, Austria; Royal Society for Protection of Nature, Thimphu, Bhutan; Max Planck Institute of Animal Behavior, Radolfzell, Germany; Institute of Zoology, Department of Ecosystem Management, Climate and Biodiversity, BOKU University, Gregor-Mendel-Strasse 33, 1180 Vienna, Austria; Natural History Museum Vienna, Ornithology, Burgring 7, 1010 Vienna, Austria

**Author notes:** Co-correspondence: Martin Kapun & Swen C. Renner.

**Keywords:** conservation genomics, effective population size, critically endangered species, Ardeidae, Himalayan riverine biodiversity

## Abstract

The White-bellied Heron (*Ardea insignis*) is one of the world’s rarest birds, with fewer than 60 known individuals remaining in the wild. Whether this extreme rarity reflects a recent anthropogenic collapse or a long history of persistently small population size has remained unknown, limiting our understanding of the species’ evolutionary resilience and conservation needs. Here, we present the first high-quality reference genome for *A. insignis*, generated using Oxford Nanopore long-read sequencing and complemented with Illumina whole-genome data. Comparative mitochondrial and nuclear phylogenomic analyses consistently recover *A. insignis* as the sister species of Purple Heron (*A. purpurea)*, while revealing moderate mitonuclear discordance among deeper ardeid lineages. Genome-wide analyses demonstrate exceptionally low heterozygosity and extensive runs of homozygosity relative to the widespread and closely related Great Blue Heron (*A. herodias*), indicating pronounced genomic erosion and long-term inbreeding. However, the predominance of short and intermediate-length homozygous tracts, together with robust Pairwise Sequentially Markovian Coalescent (PSMC) reconstructions across alternative parameterizations, indicates that *A. insignis* has persisted with comparatively small effective population sizes over much of its evolutionary history rather than experiencing only a recent demographic collapse. The two sampled individuals nevertheless differ in the abundance of longer homozygous tracts, indicating that inbreeding accumulated over the past few generations has not been uniform among the surviving birds, despite their shared history of chronic rarity. Our results indicate that the White-bellied Heron represents a lineage that has survived prolonged demographic adversity and that its greatest genetic challenge may be limited adaptive potential rather than recent genomic deterioration alone. Beyond providing the first genomic resource for this critically endangered species, our study establishes an evolutionary baseline for future monitoring and highlights the importance of integrating genomic and ecological data to guide conservation strategies for species persisting at the edge of extinction.

## Introduction

Understanding how historical and contemporary demographic processes shape genomic diversity in natural populations is a central goal in molecular ecology (Ellegren and Galtier 2016). Fluctuations in population size leave characteristic signatures across genomes that can be used to reconstruct population history over evolutionary timescales (Li and Durbin 2011), whereas habitat fragmentation and reduced connectivity structure genetic variation among contemporary populations and can create detectable barriers to gene flow (Renner et al. 2016). The increasing accessibility of whole-genome sequencing has transformed our ability to infer these processes, allowing demographic history, inbreeding, and effective population size to be estimated from genome-wide patterns of variation rather than from a limited number of molecular markers (Li and Durbin 2011; Ellegren 2014; Supple and Shapiro 2018). Such approaches have become particularly valuable in conservation biology because they provide insight into the evolutionary consequences of population decline over timescales that exceed the temporal scope of ecological monitoring (Allendorf et al. 2010; Kardos et al. 2021).

Small and isolated populations are expected to experience increased genetic drift, reduced effective population size, elevated inbreeding and a progressive loss of genetic diversity, which together erode adaptive potential and increase extinction risk (Frankham 1996; Frankham et al. 2014; Kardos et al. 2021). The genomic consequences of rarity are, however, not always straightforward. Species that have declined recently under anthropogenic pressure may still retain substantial standing genetic variation inherited from historically large populations. By contrast, species with persistently small effective population sizes for thousands of generations may show chronically low diversity. At the same time they may carry a reduced burden of strongly deleterious recessive alleles, because repeated exposure of such alleles in the homozygous state allows them to be efficiently purged, enabling some lineages to persist despite long-term inbreeding (Robinson et al. 2018; Dussex et al. 2021; Bertorelle et al. 2022). These contrasting demographic histories imply fundamentally different expectations for future evolutionary potential, genetic load and the role of genetic management. Whereas recently collapsed populations may require urgent measures to retain remaining genetic variation and mitigate inbreeding, long-term naturally rare species may benefit more from safeguarding the ecological conditions that have allowed them to persist despite chronically small population sizes. Although both scenarios demand effective habitat protection, they can require markedly different conservation strategies despite producing almost identical present-day census sizes (Hohenlohe et al. 2021). Distinguishing between them is therefore a prerequisite for evidence-based conservation, and requires genomic rather than demographic data.

Whole-genome data offer several complementary approaches for reconstructing demographic history with respect to this problem. Genome-wide heterozygosity provides a broad measure of standing genetic variation, whereas runs of homozygosity (ROH) yield a direct record of realized inbreeding (Ceballos et al. 2018; Silva et al. 2024). Because recombination progressively erodes homozygous chromosomal segments over time, the length distribution of ROH is informative about the timing of demographic events: long ROH indicate recent mating between close relatives, whereas numerous short ROH generally reflect more ancient bottlenecks or sustained small effective population sizes (Shafer et al. 2015; Ceballos et al. 2018). ROH analyses have consequently become a standard component of conservation genomics and provide substantially greater resolution than traditional pedigree-based estimates of inbreeding – an advantage that is decisive in species for which pedigrees do not exist.

Long-term demographic trajectories can be reconstructed from a single diploid genome using coalescent-based approaches such as the Pairwise Sequentially Markovian Coalescent (PSMC), which infers temporal changes in effective population size from the distribution of heterozygous sites along chromosomes (Li and Durbin 2011). PSMC has become one of the most widely applied methods for investigating long-term demographic history in both model and non-model organisms alike, including numerous threatened vertebrates (Nadachowska-Brzyska et al. 2016). Recent work has nevertheless demonstrated that commonly used default parameter settings can generate spurious peaks in effective population size followed by apparent collapses, and that these artefacts can largely be avoided by splitting the first time interval (Hilgers et al. 2025). Because demographic reconstructions are increasingly used to justify management decisions (Díez-del-Molino et al. 2018; Hohenlohe et al. 2021), validating inferences across alternative parameterizations is no longer optional, but represents a requirement to avoid spurious and biased estimates.

These analytical advances have coincided with rapid improvements in sequencing technology. Long-read sequencing platforms now enable highly contiguous reference genomes to be generated for non-model organisms from relatively small amounts of DNA (Bein et al. 2025; Schell et al. 2025), while complementary short-read data provide the sequencing accuracy required for reliable variant discovery. High-quality reference genomes have therefore become foundational resources in conservation biology: they improve read mapping, enable robust variant calling, and permit chromosome-scale analyses of structural variation, homozygosity and historical demography (Rhie et al. 2021). They also underpin comparative work, because the growing availability of avian genomes allows phylogenomic analyses based on hundreds to thousands of conserved nuclear orthologues that complement traditional mitochondrial datasets. Comparisons between mitochondrial and nuclear phylogenies are particularly informative, since discordant topologies may arise through incomplete lineage sorting, historical introgression, or differences in the evolutionary dynamics of maternally and biparentally inherited genomes (Toews and Brelsford 2012; Edwards et al. 2016; DeRaad et al. 2023; Salles et al. 2025; Zhao et al. 2025). Such mitonuclear comparisons also provide an independent assessment of assembly quality and orthologue recovery (Allio et al. 2020; Quattrini et al. 2023). These approaches are particularly valuable for threatened species, for which genomic resources are often lacking despite their importance for evolutionary and conservation research.

The White-bellied Heron (*Ardea insignis*) is among the rarest birds on Earth and provides an unusually powerful system in which to address these questions. Fewer than 60 known individuals are confirmed to survive across the riverine foothills of the eastern Himalayas in Bhutan, north-eastern India, and northern Myanmar (Kyaw et al. 2021; Royal Society For Protection of Nature 2026), and the species is classified as Critically Endangered (BirdLife International 2018; IUCN Red List 2026). Despite its evolutionary and conservation significance, genomic resources for *A. insignis* are essentially absent, reflecting a broader lack of chromosome-scale genomes across the heron family, for which assemblies are currently available for only a handful of species (Zheng et al. 2024). Within several avian familias, including Ardeidae, genomic resources remain scarce, with chromosome-scale assemblies available for only a handful of species(Zheng et al. 2024), and for critically endangered members of the family –precisely those for which sampling opportunities are most limited– they are essentially absent, such as for *A. insignis*. Ecologically it departs markedly from its congeners. Whereas most *Ardea* species exploit a broad spectrum of wetland habitats, *A. insignis* is a specialist of undisturbed, fast-flowing rivers and is confined to an interconnected network of river reaches spanning roughly 100–1,800 m a.s.l. (Kyaw et al. 2021). Within these systems it depends on a mosaic of pools, riffles, runs, side channels and ponds, taking fish – predominantly *Schizothorax*, *Garra* and related genera– by a stand-and-wait hunting technique (Khandu et al. 2021). In Bhutan the species has been recorded from all major river basins, but the majority of the known national population is concentrated in the Punatshangchhu basin.

Its breeding biology is equally restrictive. Pairs nest solitarily and at very low density on large emergent trees within well-vegetated broadleaved or chir pine forest adjacent to suitable river reaches, with neighbouring nests separated by tens of kilometres, and the breeding season in Bhutan extends from February to July (Acharja 2019; Acharja et al. 2025). Reproductive output is low, with small clutches and high mortality during the post-fledging period (Acharja 2019; Acharja et al. 2025). Movement ecology remains poorly resolved, but individuals are assumed to move seasonally along river corridors between foraging and breeding areas, so that longitudinal connectivity between reaches and the integrity of riparian vegetation are central to the persistence of the species. Besides its ecological traits, its habitat has been transformed within a few decades by hydropower development, sand and boulder extraction, riparian degradation and disturbance (Maheswaran et al. 2021). The ecological traits and the threat to its habitat highlight the need to identify the mechanism underlying the species’ present rarity.

Dependence on undisturbed, low-gradient river reaches, naturally low breeding densities and limited reproductive output are consistent with a species that has never been abundant, in which case small population size – and any associated depletion of genetic diversity – would be an ancestral condition rather than a recent one. Set against this is the accelerated transformation of these river systems over the past few decades, which offers an equally plausible mechanism for a rapid and recent decline. The two scenarios are not mutually exclusive, but they predict contrasting genomic signatures – uniformly low diversity dominated by short ROH under chronic rarity, as against retained diversity punctuated by long ROH under recent collapse – while remaining indistinguishable from census data alone.

To date, the only genomic resource published for *A. insignis* is a complete mitochondrial genome (Duan et al. 2018); a nuclear reference genome is lacking entirely. This gap constrains several questions that are fundamental to both evolutionary biology and conservation. First, it remains unclear whether the present-day rarity of the species primarily reflects a recent anthropogenic decline or the persistence of small effective population sizes throughout much of its evolutionary history. Second, the extent of genome-wide inbreeding and the loss of genetic diversity are unknown. Third, without a high-quality reference genome, comparative evolutionary analyses and future population genomic work – including any genetically informed management of the captive population – remain out of reach. Resolving these questions matters because the timing and magnitude of demographic decline directly determine expectations regarding adaptive potential, genetic load and the likely effectiveness of conservation intervention.

Here we present the first high-quality reference genome for the *A. insignis* generated using Oxford Nanopore long-read sequencing in combination with Illumina short-read data. We combine de novo genome assembly, chromosome-scale scaffolding, comparative mitochondrial and nuclear phylogenomics, and population genomic analyses to investigate the evolutionary history of this critically endangered species. Specifically, we (i) generate and evaluate a highly contiguous genome assembly, (ii) infer the phylogenetic placement of *A. insignis* within the family Ardeidae using both mitochondrial genomes and hundreds of conserved nuclear BUSCO orthologues, (iii) quantify genome-wide homozygosity and inbreeding using runs of homozygosity, and (iv) reconstruct long-term demographic history using PSMC while explicitly accounting for recently identified methodological artefacts associated with standard PSMC parameterization. Together, these analyses provide the first genome-scale perspective on the evolutionary history of one of the world’s rarest birds and establish a genomic foundation for future ecological, evolutionary, and conservation research on *A. insignis*.

## Materials and Methods

### Sampling, DNA extraction and sequencing

Population genomic analyses were based on two White-bellied Herons (*Ardea insignis*) sampled in Bhutan between 2025 and 2026 (Figure 1 and Table 1). One individual (“Wing”) was sampled in March 2026. This bird, which originated from the wild at the Lhamoizingkha (Dagana), downstream of Punatshangchhu, has an amputated wing and is held in captivity at the White-bellied Heron Conservation Center (WBHCC). A second individual (“Wire”), which had died following collision with electric wires near Tingtibi, Mangdechhu, in July 2025, was sampled alive in May 2025 prior to its death (Figure 1 and Table 1). Blood samples were preserved in 70% lab grade ethanol until DNA extraction.

**Figure 1.**
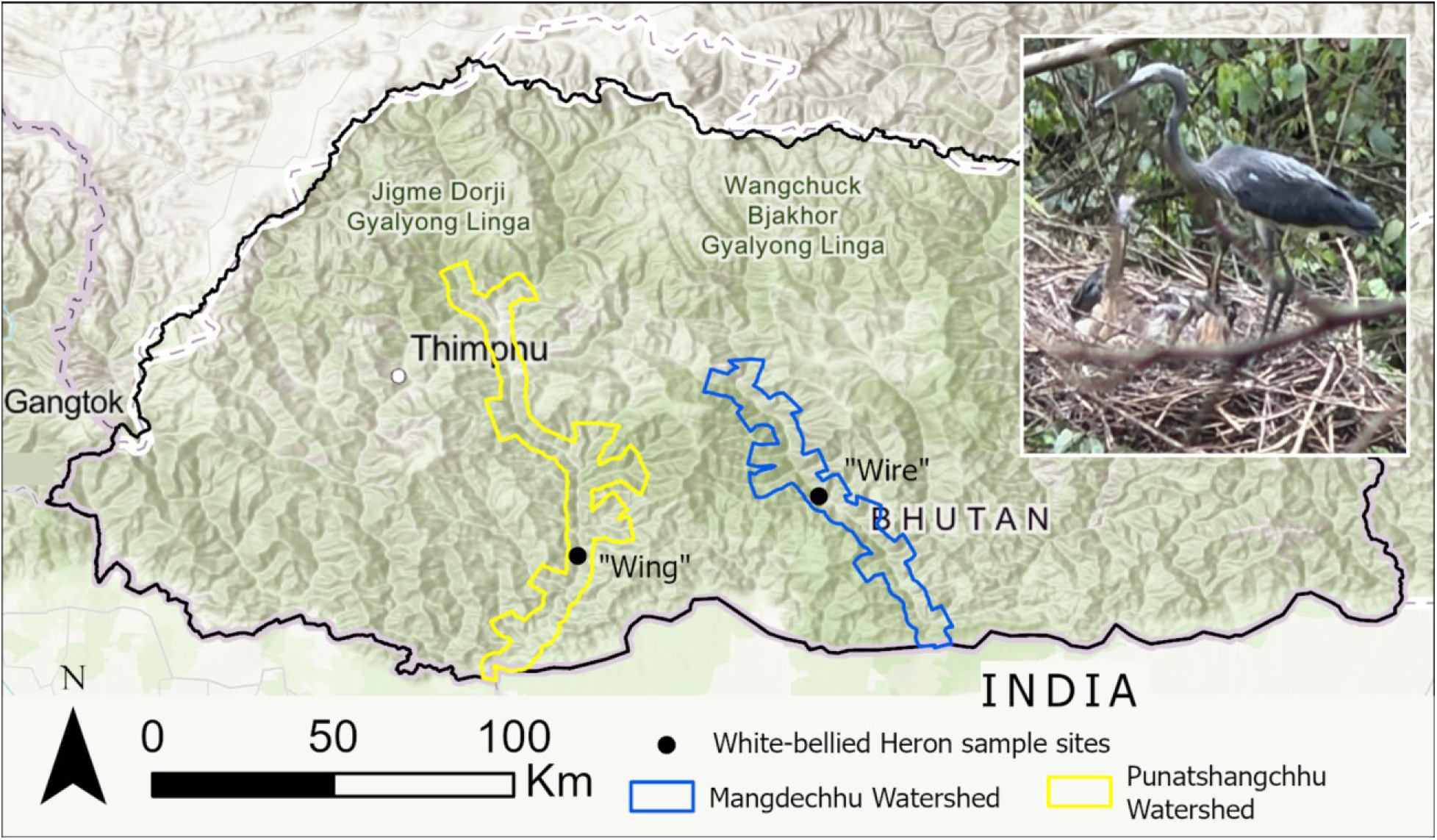
**Sampling sites and distribution of *A. insignis* specimens analyzed in this study.**

**Table 1.**
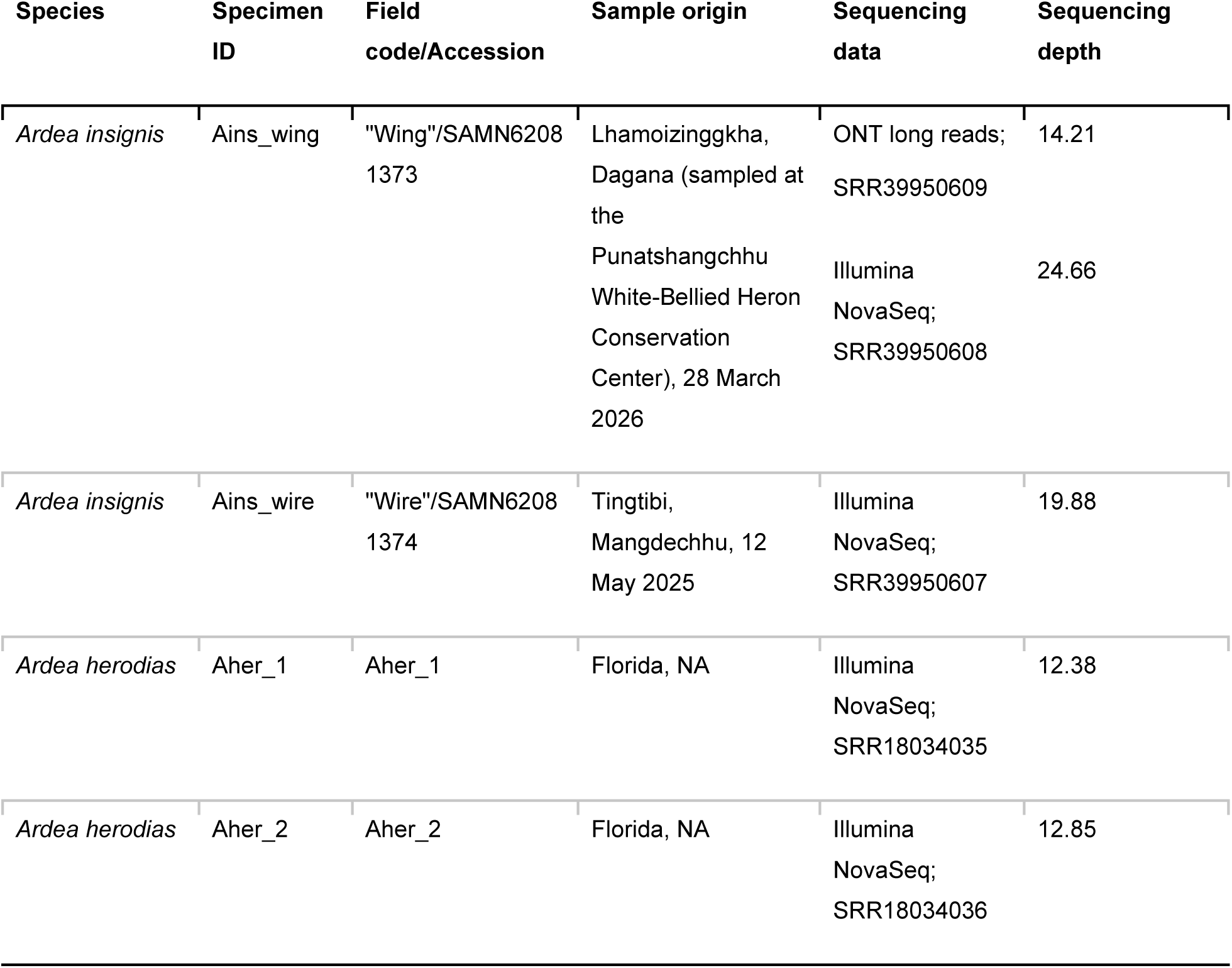
Samples included in the population genomic analyses. , with specimen identifiers, sampling information, sequencing data, and mean genome-wide sequencing depth.

Genomic DNA was extracted using the QIAwave DNA Blood & Tissue Kit (QIAGEN) following the manufacturer’s protocol with the following modifications: incubation at 56 °C for 1 h, elution in 70 µl NFW. DNA concentration was quantified using the QuantiFluor ONE dsDNA System (PROMEGA) and fragment size distributions were assessed using the Genomic DNA Screen Tape analysis (Agilent).

To generate a reference genome, high-molecular-weight DNA from “Wing” was sequenced on an Oxford Nanopore MinION Mk1C platform using an R10.4.1 flow cell and the SQK-NBD114-24 library preparation kit. Raw electrical signals were base-called with Dorado (v.1.4.0; Oxford Nanopore Technologies 2026) using the super-accuracy model, including adapter trimming and barcode demultiplexing. Read quality and length distributions were assessed with NanoPlot (De Coster et al. 2018). Complementary Illumina paired-end sequencing was performed by Macrogen Europe B.V. (Amsterdam, Netherlands) for both individuals after paired-end library preparation using the NEBNext® Ultra II FS DNA Library Prep kit for Illumina, with the following specifications: protocol for use with inputs > 100 ng, fragmentation time 8 min, bead based size selection aiming for an insert size distribution of 275 to 475 bp. Illumina indices used for multiplexed sequencing are available in Table S1. The resulting paired-end reads of 150 bp lengths were used for assembly validation and downstream population genomic analyses. Raw reads were deposited in the NCBI Short Read Archive under BioProject PRJNA1505165)

### Genome assembly and annotation

We first estimated genome characteristics from the Oxford Nanopore reads using GenomeScope 2.0 (Ranallo-Benavidez et al. 2020) to infer haploid genome size, repeat content, and heterozygosity from the *k*-mer frequency distribution. Subsequently, we performed de-novo assembly of the reference genome based on untrimmed reads using Flye v2.9, following the developers recommendations and employing the Oxford Nanopore high-quality mode (Kolmogorov et al. 2019). The initial assembly was subsequently polished through four iterative rounds of Racon (v.1.5.0; Vaser et al. 2017), with Illumina reads from the same specimen, that were initially trimmed with FASTP (v.1.3.3; Chen et al. 2018) and then mapped back to the assembly using minimap2 (v.2.28; Li 2018) before each polishing step.

Assembly quality was assessed using several complementary approaches. Assembly contiguity statistics, including assembly size, N50, L50, and contig number, were calculated with QUAST (Gurevich et al. 2013). Gene-space completeness was evaluated with BUSCO (v.6.0.0; Manni et al. 2021) using the *vertebrata_odb10* lineage dataset (Kriventseva et al. 2019) and the MetaEuk gene predictor (v.7.bba0d80; Levy Karin et al. 2020). To identify potential contaminant contigs, Oxford Nanopore reads were mapped back to the polished assembly with minimap2 and contigs were taxonomically classified using BLAST against the NCBI nucleotide database. Coverage, GC content, and taxonomic assignments were subsequently integrated using BlobTools2 (Laetsch and Blaxter 2017).

To facilitate chromosome-scale analyses, the polished assembly was scaffolded using RagTag (v.2.1.0; Alonge et al. 2022) with the chromosome-level genome assembly of the Chinese Egret (*Egretta eulophotes*; GCA_056459225.1) as the reference. Contigs lacking unambiguous alignments were retained as unplaced scaffolds. Synteny between the *A. insignis* assembly and the reference genome was evaluated using whole-genome alignments generated with minimap2 and visualized as pairwise alignment dot plots in R using ggplot2 (Wickham et al. 2019).

The mitochondrial genome of *A. insignis* was extracted directly from the assembled genome using MitoFinder (v.1.4.1; Allio et al. 2020), employing a combination of 21 mitochondrial reference genomes available for the Ardeidae family from the NCBI RefSeq database as the annotation reference. Assembly and annotation were further refined using a combined mapping and re-assembly approach and manual verification of MITOS2 (v.2.1.10; Bernt et al. 2013; Donath et al. 2019) annotations (see the Supplementary Materials for details). Annotated mitogenome contigs from previously available genomes identified by MitoFinder were deposited in GenBank format at the NHM data repository (Kapun et al. 2026).

### Comparative phylogenomic analyses

To determine the phylogenetic placement of *A. insignis* within the family Ardeidae, we reconstructed both mitochondrial and nuclear phylogenies. Throughout the analyses, taxon names and species delimitation followed the avian taxonomy of Clements et al. (2025). Twenty-one publicly available Ardeidae mitochondrial genomes, including one of *A. insignis*, together with the outgroup *Ciconia ciconia* were downloaded from NCBI RefSeq (Table S2). We complemented and extended this dataset by reconstructing five additional complete mitochondrial genomes based on available Ardeidae reference genomes that were downloaded from the RefSeq database (Table S2) using MitoFinder in assembly search mode (−o 5) (Allio et al. 2020), applying a minimum contig length threshold of 150 bp. MitoFinder was further used to annotate mitochondrial genes, including the 13 protein-coding genes, two ribosomal RNAs, and 22 transfer RNAs characteristic of our de novo assembly of the “Wing” specimen as well as the other avian mitochondrial genomes. We additionally used MitoFinder in de-novo assembly mode (-o 2) to reconstruct mitochondrial genome from raw reads of the *A. herodias* sample Aher_1 (Table 1) using the MegaHit assembler (Li et al. 2015) with parameter specific to MitoFinder. Annotation followed the same approach as for the novel *A. insignis* mitogenome detailed in the Supplementary Materials.

First, we focused on the 13 mitochondrial protein-coding genes and two rRNA genes, which were extracted from the annotated GenBank files. Each gene was aligned separately across all individuals before being concatenated into a partitioned supermatrix using a custom Python script. Maximum-likelihood phylogenetic inference was then performed in IQ-TREE3 (v.3.0.1; Minh et al. 2020) using the partition-merging strategy implemented in ModelFinder (Kalyaanamoorthy et al. 2017), with branch support assessed by 1,000 ultrafast bootstrap replicates (Hoang et al. 2018).

To complement the mitochondrial analyses, 15 chromosome-and scaffold-level genome assemblies publicly available for Ardeidae were downloaded from NCBI (Table S3). Conserved single-copy orthologues were identified using BUSCO with the aves_odb10 database in MetaEuk mode, yielding amino acid and nucleotide coding sequences, respectively. To further include the genomic information of the second *A. insignis* specimen (“Wire”) into the analysis, we mapped the trimmed Illumina reads against the BUSCO sequences of the “Wing” individual with minimap2 and used samtools consensus to reconstruct consensus sequences that were then integrated in the BUSCO dataset. Then, we aligned BUSCO genes with MAFFT, we filtered for genes that were (1) present in all samples, (2) whose alignment was larger than 800 bp and (3) which contained less than 10% missing data for downstream analysis. The retained loci were concatenated into a partitioned supermatrix, and a maximum-likelihood species tree was inferred in IQ-TREE using the same partition-merging strategy and branch-support assessment described for the mitochondrial protein-coding gene analysis.

### Inference of inbreeding and genome-wide levels of heterozygosity

We first trimmed and cleaned raw Illumina reads of both individuals from adapter sequences with FASTP Then, we mapped the trimmed reads against the *A. insignis* reference genome, scaffolded with RagTag to pseudo-chromosome-level, using minimap2 with default parameters. Finally we generated duplicate-aware, coordinate-sorted alignments for downstream analyses using the samtools (v.1.18; Danecek et al. 2021) modules *markdup* and *sort*, respectively.

Single nucleotide variants were called for both samples jointly using BCFtools (v.1.16; Li 2011) with minimum mapping and base quality thresholds of 20. To minimize biases caused by low coverage and repetitive regions, depth filters were calculated individually for each sample, excluding sites with coverage below one-third or above twice the mean sequencing depth. We subsequently counted the number of genome-wide heterozygous sites for each sample to estimate specimen-specific amounts of genetic variation.

Complementary to this, runs of homozygosity (ROHs) were identified using the hidden Markov model implemented in bcftools roh (Narasimhan et al. 2016). Because population allele frequency estimates were unavailable for *A. insignis*, a constant default alternative allele frequency of 0.5 (--AF-dflt 0.5) was specified. This value corresponds to a non-informative prior that assumes maximal expected heterozygosity and avoids introducing bias towards either common or rare alleles in the absence of empirical allele frequency information. ROHs were inferred from genotype likelihoods across all callable sites, including invariant positions, using the default HMM parameters of bcftools roh and then classified into four length classes (100 kb– 1 Mb, 1–5 Mb, 5–10 Mb, and >10 Mb), allowing recent and more ancient inbreeding to be distinguished (Pemberton et al. 2012; Ceballos et al. 2018). Individual genomic inbreeding coefficients were estimated as the proportion of the assembled genome contained within ROHs.

To place the genomic diversity and inbreeding patterns of *A. insignis* into a broader evolutionary context, we analysed two publicly available whole-genome datasets of the widespread Great Blue Heron (*Ardea herodias*; GenBank SRA accessions SRR18034035 and SRR18034036; Table 1) using the identical analytical pipeline. In the absence of a chromosome-scale genome assembly from a more closely related *Ardea* species, reads from both species were mapped against the *A. insignis* reference genome. Mapping the more divergent *A. herodias* reads to an *A. insignis* reference may reduce mapping efficiency at highly divergent loci, thereby leading to conservative estimates of heterozygosity and potentially inflated ROH lengths in *A. herodias*. Consequently, any reduction in genomic diversity observed in *A. insignis* is likely conservative, as reference bias would tend to underestimate diversity in *A. herodias* rather than artificially inflate diversity in *A. insignis*.

### Historical demographic inference

Historical changes in effective population size (*N*e) were reconstructed using the Pairwise Sequentially Markovian Coalescent (PSMC v.0.6.5; Li and Durbin 2011). Similar to above, we performed these analyses for both the two *A. insignis* and the two *A. herodias* specimens to compare the demographic history of the critically endangered White-bellied heron with that of a widespread congener. Using the coordinate-sorted BAM files generated above, we produced diploid consensus sequences with bcftools applying the same minimum mapping and base quality thresholds of 20. Sample-specific depth filters corresponding to one-third and twice the mean sequencing depth were applied prior to conversion into PSMC input format, thereby excluding poorly supported and excessively high-coverage regions. We further masked repetitive regions and restricted our analyses to the RagTag pseudochromosomes to minimize mapping artefacts and avoid biases arising from fragmented or unplaced scaffolds.

To account for recently identified artefacts associated with the default PSMC time-interval scheme, particularly spurious recent effective population size peaks caused by grouping the first four atomic intervals into a single broad time bin (Hilgers et al. 2025), demographic histories were reconstructed using three alternative interval parameterizations: the default setting (4+25*2+4+6), a two-way split of the most recent interval (2+2+25*2+4+6), and a four-way split of this interval (1+1+1+1+25*2+4+6). The latter two parameterizations provide increased resolution in the most recent part of the trajectory and were used to assess whether inferred demographic features were robust to the interval scheme. Demographic patterns were considered robust only when consistently recovered across the modified parameterizations, whereas features restricted to the default setting were interpreted cautiously as potential methodological artefacts. Statistical confidence was assessed using 100 bootstrap replicates per individual and parameterization. Demographic trajectories were scaled assuming a generation time of 10 years and a neutral mutation rate of 1.23 × 10^-9^ substitutions per site per year (Nadachowska-Brzyska et al. 2016).

## Results

### Genome assembly provides a highly contiguous and complete reference genome for *Ardea insignis*

Oxford Nanopore sequencing of the *A. insignis* individual “Wing” generated 3,531,001 reads comprising 19.3 Gb of sequence data, corresponding to approximately 14.21x genome coverage (Table 1). Sequencing quality was high, with a median read quality of Q24.5; 93.7% of reads exceeded Q10 and 78.3% exceeded Q20. The read-length N50 reached 9,445 bp, while the longest read measured 1,256,892 bp, matching TapeStation measurements and confirming the availability of high-molecular-weight DNA suitable for long-read genome assembly.

GenomeScope analysis of the *k*-mer frequency spectrum estimated a haploid genome size of 1.21 Gb (1.207–1.215 Gb), with an estimated repeat content of 100.9–101.6 Mb (approximately 8.4% of the genome). The model fit to the observed *k*-mer distribution ranged between 93.1% and 99.7%, indicating an excellent fit of the GenomeScope model and a single dominant homozygous peak without evidence of pronounced haplotype separation.

De-novo assembly yielded a draft genome of 1,198,150,322 bp length, distributed across 1,671 contigs, with a contig N50 of 4.64 Mb, a contig N90 of 975 kb and a maximum contig length of 15.0 Mb (Figure 2A). The assembled genome exhibited a GC content of 43.2% and closely matched the expected genome-size estimated with GenomeScope, indicating near-complete recovery of the genome. BUSCO analysis recovered 95.3% of the 3,354 conserved vertebrate single-copy orthologues, while only 2.9% were classified as missing. Duplicated BUSCOs accounted for only 0.5% of the dataset, indicating efficient collapse of alternative haplotypes during assembly. The BlobTools analyses of the draft genome detected no evidence of substantial contamination, with virtually all contigs assigned to Aves and exhibiting coverage and GC-content profiles consistent with a single avian genome (Figure 2B).

**Figure 2.**
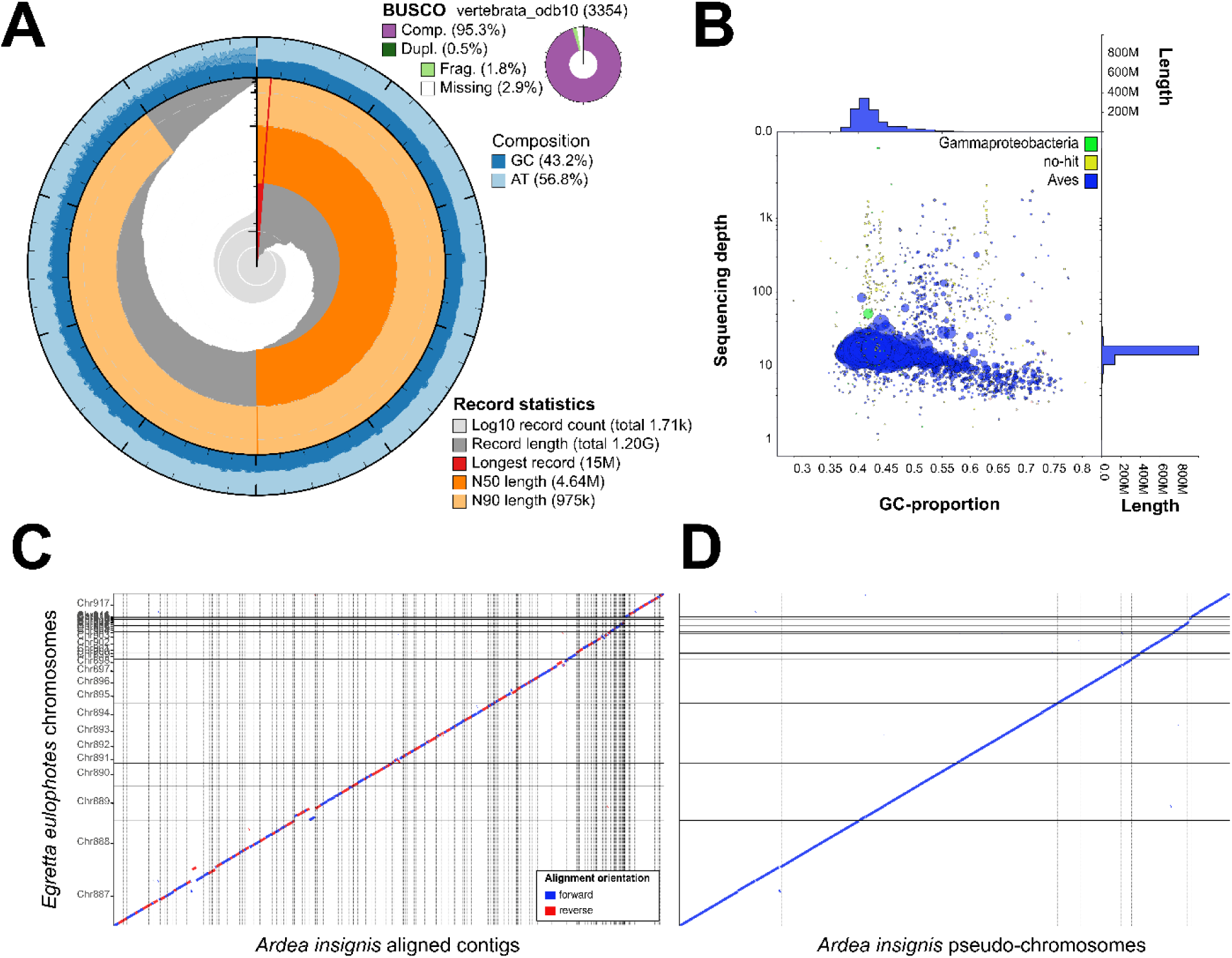
Genome assembly quality assessment and chromosome-scale scaffolding of the *Ardea insignis* reference genome. (A) BlobTools snail plot summarizing assembly completeness and quality. Concentric tracks show scaffold length distribution, GC content, BUSCO completeness (vertebrata_odb10), and the proportion of assembled sequence assigned to major taxonomic groups. The assembly comprises approximately 1.20 Gb with 95.3% complete BUSCO genes and is largely assigned to avian sequences. (B) BlobTools GC-content versus sequencing coverage plot showing the taxonomic assignment of assembled contigs. Circle size is proportional to contig length, and colours indicate taxonomic classification based on the best BLAST hit against the NCBI nucleotide database. The assembly is dominated by contigs assigned to Aves, with no evidence of substantial non-avian contamination. Marginal histograms summarize the distribution of GC content and sequencing coverage. (C) Whole-genome alignment of the raw *A. insignis* contigs against the chromosome-level genome of the Chinese Egret (*Egretta eulophotes*) prior to scaffolding. Forward and reverse alignments are shown in blue and red, respectively, revealing extensive chromosome-scale synteny despite fragmentation of the initial assembly. (D) Whole-genome alignment following reference-guided scaffolding with RagTag. The nearly continuous diagonal indicates successful ordering and orientation of contigs into chromosome-scale pseudomolecules and demonstrates high collinearity between the *A. insignis* genome and the *E. eulophotes* reference, with only a small number of minor rearrangements or unplaced scaffolds remaining.

Reference-guided scaffolding against the chromosome-level reference genome of *Egretta eulophotes* anchored 1,075 contigs, representing 99.5% of the assembled sequence, onto chromosome-scale pseudochromosomes, leaving only 0.5% of the assembly unplaced (Figure 2C). Whole-genome alignments demonstrated extensive chromosome-scale collinearity between *A. insignis* and *E. eulophotes*, with no evidence for major chromosomal rearrangements (Figure 2D). Together, these results demonstrate that the assembled genome is highly contiguous, complete and suitable for downstream comparative phylogenomic and population genomic analyses.

### Comparative phylogenomics places *Ardea insignis* within the large-heron radiation

We successfully recovered the complete mitochondrial genome from the *A. insignis* de novo assembly and analyzed the mitochondrial sequences together with publicly available Ardeidae mitogenomes. Phylogenomic analyses based on partitioned alignments of protein-coding and rRNA genes successfully recovered highly congruent topologies. The newly assembled mitochondrial genome clustered with the previously published *A. insignis* mitogenome with maximal support (UFBoot = 100), confirming the accuracy of the assembly. Within the genus *Ardea*, *A. insignis* was consistently recovered as sister to *A. purpurea* with moderate bootstrapping support (UFBoot = 87; Figure 3A).

**Figure 3.**
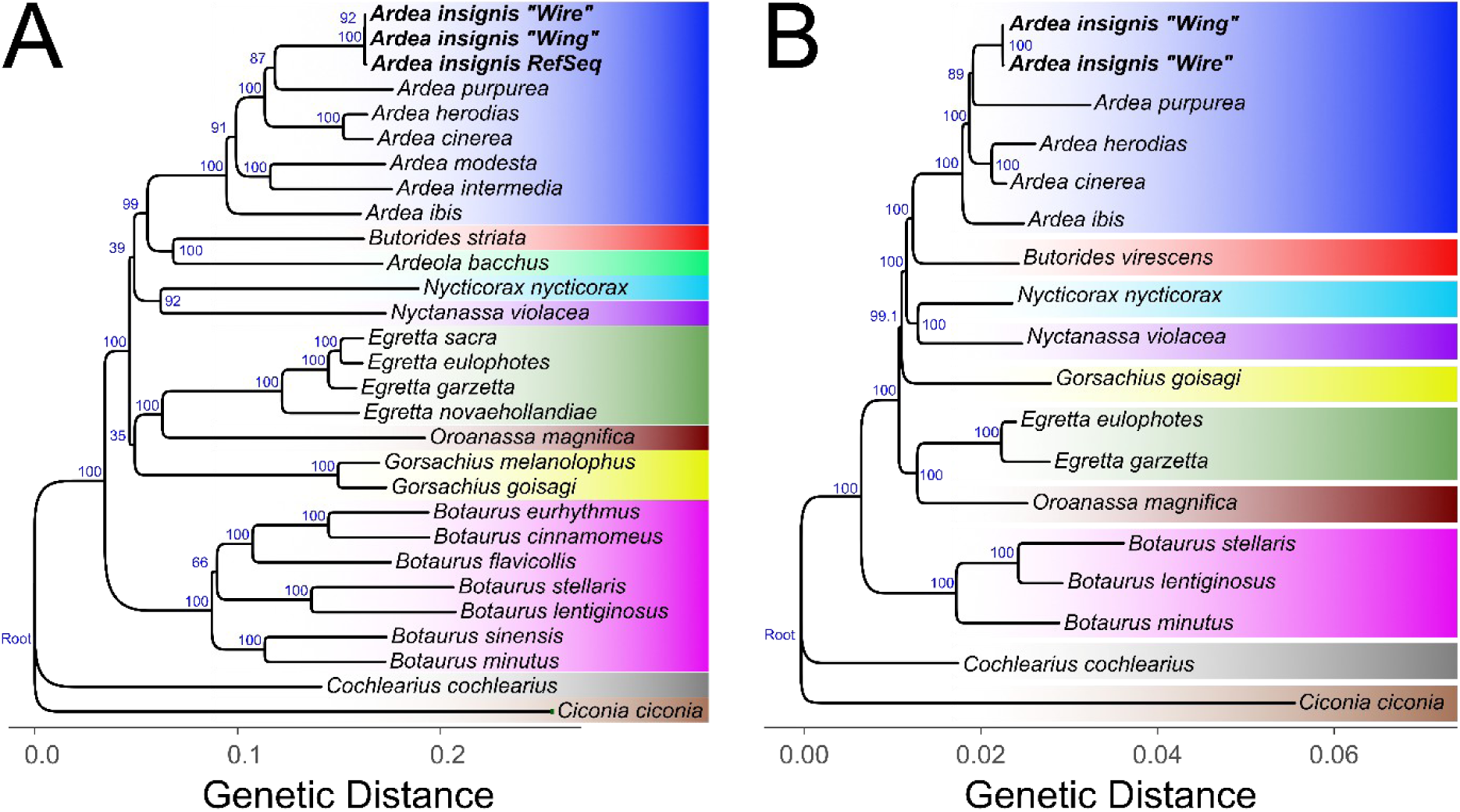
Phylogenetic placement of the White-bellied Heron (*Ardea insignis*) inferred from mitochondrial and nuclear genomic data. (A) Maximum-likelihood phylogeny reconstructed from a partitioned supermatrix comprising all 13 mitochondrial protein-coding and two rRNA genes. (B) Maximum-likelihood phylogeny inferred from a concatenated alignment of 573 conserved single-copy nuclear BUSCO genes that fulfilled the following criteria: (1) presence in all analyzed taxa, (2) alignment length >800 bp, and (3) <10% missing data across the alignment. Branch support was assessed using ultrafast bootstrap approximation (UFBoot2; 1,000 replicates). Support values are shown at the nodes as UFBoot2 percentages. The newly assembled *Ardea insignis* genomes of “Wing” and “Wire” are highlighted in bold. Both mitochondrial and nuclear phylogenies consistently recover *A. insignis* as sister to *A. purpurea* within the genus *Ardea* with moderate statistical support. *Ciconia ciconia* was used as the outgroup to root both phylogenies. Branch lengths are proportional to the number of substitutions per site, and colored bars indicate the major lineages within the family Ardeidae.

To compare mitochondrial and nuclear evolutionary histories, phylogenetic relationships were additionally reconstructed from a concatenated dataset of 573 conserved BUSCO orthologues. Consistent with the mitogenomic analyses, the nuclear nucleotide supermatrix recovered *A. insignis* as the sister species of *A. purpurea* with similarly moderate support (UFBoot = 89; Figure 3B). Overall, mitochondrial and nuclear phylogenies were highly congruent, consistently recovering the monophyly of the genera *Ardea*, *Egretta*, and *Botaurus*, as well as the sister relationship between *A. insignis* and *A. purpurea* (Figure 3). Consistent with the results from Hruska et al. (2023), both datasets recovered *Oroanassa magnifica* as the sister lineage to the monophyletic *Egretta* clade.

However, the mitochondrial and nuclear phylogenies differed in the deeper relationships among the main ardeid lineages. Both datasets consistently recovered the major genera (*Ardea*, *Butorides*, *Nycticorax*, *Nyctanassa*, *Egretta*, *Gorsachius*, *Botaurus*, and *Oroanassa*) as distinct, well-supported lineages, but their relationships differed among the non-*Ardea* taxa. In the mitochondrial phylogeny, *Ardeola* grouped with *Nycticorax*, whereas *Egretta* formed a clade sister to *Oroanassa*, which together were sister taxa to *Gorsachius*. By contrast, the nuclear BUSCO phylogeny placed *Butorides*, *Nycticorax*, *Nyctanassa*, *Gorsachius*, *Egretta*, and *Oroanassa* as successive lineages outside *Ardea*, with *Ardeola* not represented in the nuclear dataset. Thus, while the placement of *A. insignis* as sister to *A. purpurea* and the relationships within *Ardea* were fully congruent between datasets, the branching order among several deeper ardeid lineages exhibited moderate mitonuclear discordance. Importantly, most conflicting backbone nodes in the mitochondrial phylogeny received only low to very low bootstrap support, indicating limited resolution of these deeper relationships with mitochondrial markers.

### Genome-wide diversity is markedly reduced in *Ardea insignis*

While the preceding analyses resolved the evolutionary position of *A. insignis* among extant herons, the availability of a high-quality reference genome also enabled us to investigate patterns of genome-wide variation within the species. To this end, we mapped high-quality Illumina reads from two *A. insignis* individuals and two *A. herodias* individuals to the newly assembled *A. insignis* reference genome at pseudo-chromosome-level to characterize genome-wide patterns of genetic variation. After applying identical filtering criteria, we identified 112,696 high-confidence SNPs in the reference individual (“Wing”) and 377,237 SNPs in the second *A. insignis* individual (“Wire”). As expected for the individual from which the reference genome was assembled, nearly all variants detected in “Wing” represented heterozygous sites, whereas the higher SNP count in “Wire” additionally reflected fixed differences relative to the reference genome.

Across approximately 1.16 billion callable genomic sites in the two *A. insignis* individuals and 1.12 billion in the two *A. herodias* outgroups, the number of heterozygous genotype calls differed markedly between species. “Wing” contained 109,729 heterozygous sites (0.00945% of callable sites) and “Wire” contained 311,460 heterozygous sites (0.02685%), whereas the two *A. herodias* individuals harboured 1,395,593 (Aher_1; 0.12506%) and 1,351,739 (Aher_2; 0.12097%) heterozygous sites, respectively. Genome-wide heterozygosity was therefore approximately 4.5-to 13-fold higher in *A. herodias* compared to *A. insignis*, confirming an exceptionally depauperate pool of segregating variation in the endangered species.

Runs of homozygosity (ROH) inferred with HMM implemented in bcftools roh revealed pronounced and concordant autozygosity in both *A. insignis* individuals (Figure 4). “Wing” harboured 403 ROH exceeding 100 kb, spanning 337.3 Mb of the assembled genome (288 tracts of 100 kb-1 Mb with a total 127.9 Mb lengths, and 115 tracts of 1-5 Mb length with a total length of 209.3 Mb), corresponding to a genomic inbreeding coefficient of FROH = 0.28. “Wire” exhibited a qualitatively similar but somewhat less extreme pattern, with 450 ROH spanning 212.9 Mb (402 tracts of 100 kb–1 Mb totalling 143.1 Mb, and 48 tracts of 1–5 Mb totalling 69.7 Mb; FROH = 0.18). No ROH exceeded 5 Mb in either individual (Figure 4). Both *A. insignis* individuals, including the reference individual “Wing”, thus show clear and consistent evidence of substantial autozygosity.

**Figure 4.**
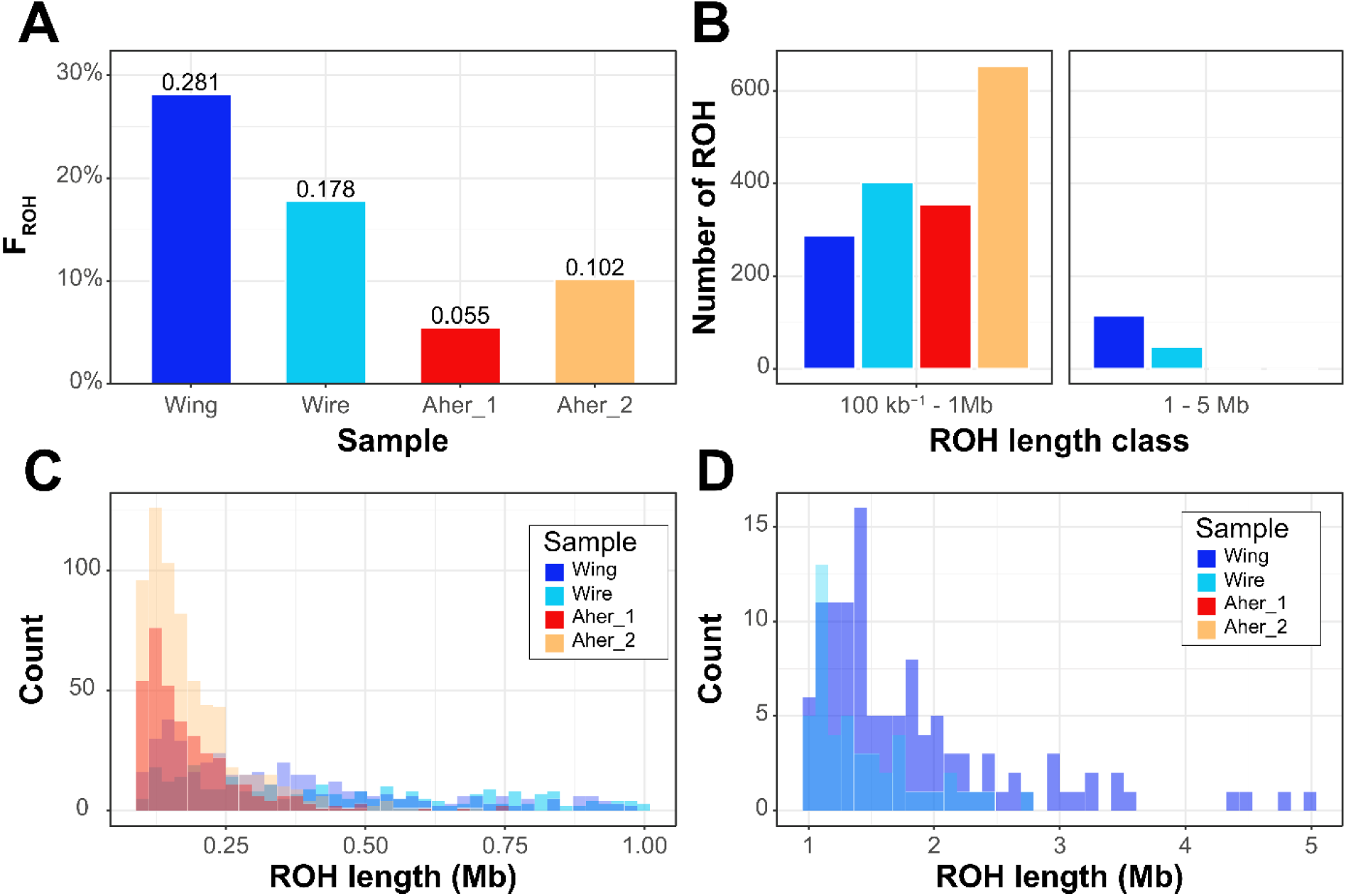
Runs of homozygosity (ROHs) reveal contrasting genomic patterns between *Ardea insignis* and *Ardea herodias*. **(A)** Fraction of the genome contained within ROHs (FROH) for each individual, calculated as the cumulative length of all ROHs ≥100 kb divided by the assembled genome size (1,198.2 Mb). Values above bars indicate the corresponding FROH. **(B)** Number of ROHs within the two principal length classes (100 kb–1 Mb and 1–5 Mb). **(C–D)** Length-frequency distributions of individual ROHs for the two length classes shown in (B), with panel **C** displaying ROHs between 100 kb and 1 Mb and panel **D** displaying ROHs between 1 and 5 Mb. Bars represent counts of ROHs within consecutive length bins for each individual. The two *A. insignis* genomes exhibit substantially greater genomic homozygosity than the two *A. herodias* genomes, reflected by higher FROH values and a marked excess of short-and intermediate-length ROHs, consistent with long-term small effective population size and persistent inbreeding rather than very recent consanguinity. In contrast, *A. herodias* individuals contain relatively few ROHs, indicating a more outbred demographic history.

In contrast, both *A. herodias* outgroup individuals contained comparatively few and exclusively short ROH. Aher_1 harboured 354 tracts between 100 kb and 1 Mb (65.8 Mb total; FROH = 0.055), and Aher_2 harboured 654 such tracts (121.7 Mb total; FROH = 0.10), with no ROH exceeding 1 Mb detected in either individual (Figure 4A-B). Moreover, within the 100 kb–1 Mb size class shared by both species, ROHs in *A. herodias* were strongly skewed toward shorter tracts of less than 500 kb, whereas those in *A. insignis* were distributed more evenly across the entire size range (Figure 4C). Such a skewed distribution pattern and the absence of long ROH together with markedly lower FROH values in the outgroup is consistent with the larger, more diverse and outbred population history typical of the widespread *A. herodias*.

Taken together, genome-wide heterozygosity and FROH consistently demonstrate that *A. insignis* possesses markedly reduced genomic diversity and elevated autozygosity relative to its congener. Our analysis shows that both sequenced *A. insignis* individuals carry a substantial fraction of their genome (18–28%) in runs of homozygosity, indicating a shared demographic history of small effective population size and/or historical inbreeding rather than an individual-specific artefact.

### Comparative demographic analyses reveal contrasting population histories of

### *Ardea insignis* and *Ardea herodias*

Historical demographic trajectories were reconstructed independently for the two *A. insignis* individuals and two publicly available *A. herodias* genomes using PSMC under three alternative time-interval parameterizations (Figure S1). Within each species, replicate analyses recovered nearly identical demographic trajectories across all parameterizations, demonstrating that the inferred demographic histories were highly reproducible despite differences in sequencing datasets and individual genomic backgrounds. Likewise, all three parameterizations produced virtually indistinguishable trajectories, indicating that the inferred demographic patterns are robust to the choice of PSMC interval scheme.

The two *A. insignis* individuals exhibited remarkably similar demographic histories. Despite pronounced differences in recent inbreeding, with one individual exhibiting substantially longer runs of homozygosity than the other (see above), both *A. insignis* genomes recovered nearly identical demographic trajectories (Figure 5). This demonstrates that the inferred demographic history reflects long-term changes in effective population size rather than recent, individual-specific inbreeding events.

**Figure 5.**
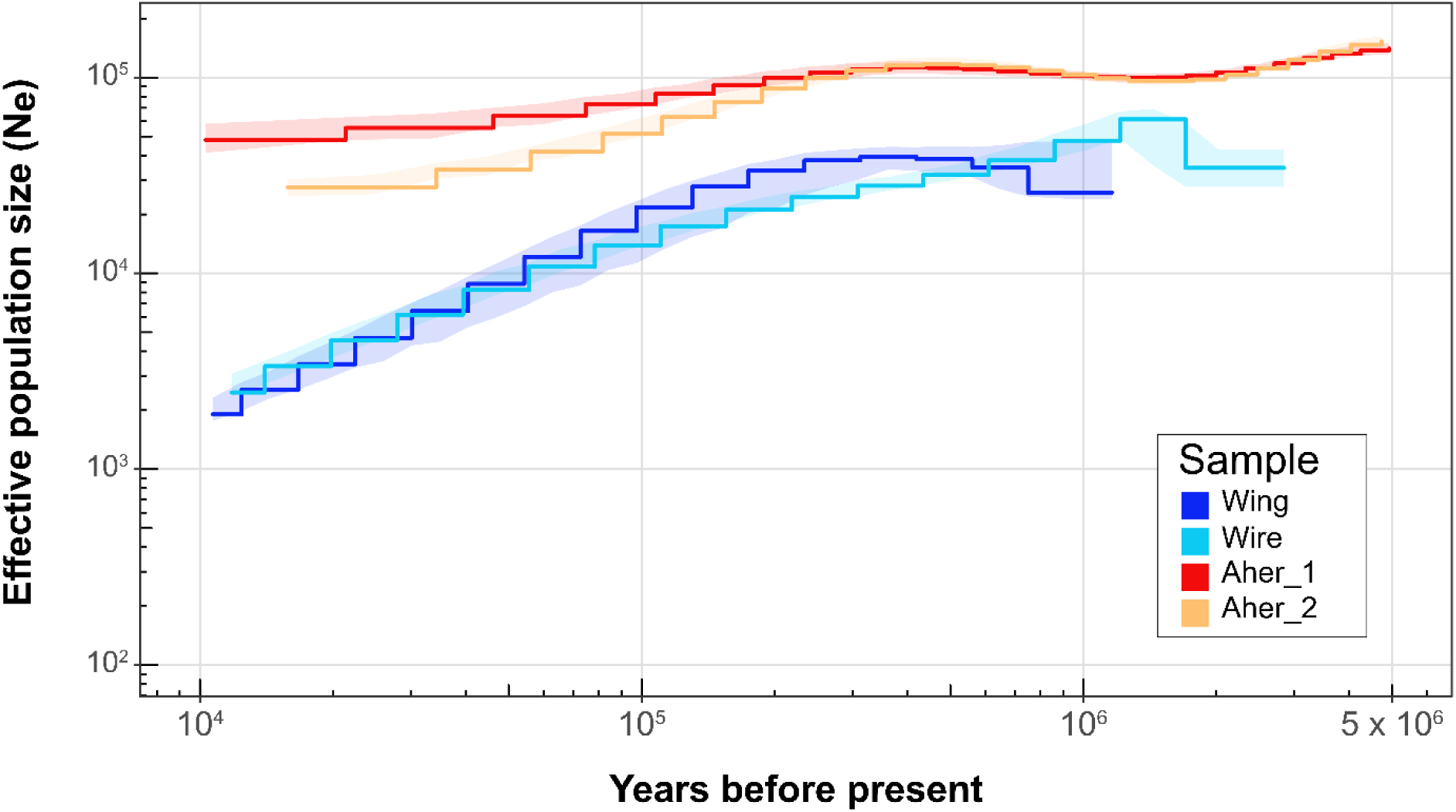
Pairwise sequentially Markovian coalescent (PSMC) estimates of historical effective population size in *Ardea insignis* and *Ardea herodias*. Historical trajectories of effective population size (Nₑ) through time are shown for two *A. insignis* individuals (“Wing” and “Wire”) and two *A. herodias* individuals (Aher_1 and Aher_2). Solid stepwise lines represent the PSMC estimates for each genome, while the corresponding shaded areas depict 95% bootstrap confidence intervals. Both axes are shown on logarithmic scales. The two *A. insignis* genomes consistently recover substantially smaller historical effective population sizes than the *A. herodias* genomes across most of the reconstructed time interval. The broadly concordant trajectories obtained from independently sequenced individuals within each species indicate that the inferred patterns primarily reflect species-level demographic histories rather than genome-specific variation. PSMC estimates toward the most recent and oldest parts of the trajectories should be interpreted cautiously because resolution is reduced at the temporal limits of the method.

Comparison with *A. herodias* revealed striking differences in long-term effective population size. Across the entire time period reconstructed by PSMC, *A. insignis* consistently exhibited substantially lower inferred effective population sizes than *A. herodias*, whose two individuals likewise showed highly concordant demographic trajectories. Although absolute estimates of effective population size should be interpreted cautiously because the genomes differed in sequencing depth, sequencing chemistry and mapping reference, the relative separation between species remained highly consistent across all parameterizations and filtering strategies (Figure S1).

Rather than indicating a recent demographic collapse, the PSMC trajectories suggest that *A. insignis* has persisted with comparatively small effective population sizes throughout much of its evolutionary history. In contrast, *A. herodias* maintained consistently much larger effective population sizes across the same time period. The close agreement between replicate individuals and across alternative PSMC parameterizations indicates that these contrasting demographic histories are robust and unlikely to represent methodological artefacts.

## Discussion

The White-bellied Heron (*Ardea insignis*) is one of the most threatened birds in the world, yet its evolutionary history and genomic status have remained largely unexplored. By combining a high-quality reference genome with comparative phylogenomics and population genomic analyses, our study provides the first genome-wide perspective on the demographic history of this critically endangered species. Three principal findings emerge: First, the newly assembled genome represents a highly complete genomic resource that allows reconstructing the phylogenetic placement of *A. insignis* within the Ardeidae family and establishes a foundation for future ecological and evolutionary studies. Second, both demographic reconstruction and genome-wide patterns of homozygosity indicate that *A. insignis* has experienced a prolonged history of small effective population size rather than solely a recent demographic collapse.

Finally, despite this long history of rarity, the genomic data suggest that the species has persisted over evolutionary timescales at small population sizes, emphasizing that extremely small populations are not necessarily evolutionarily doomed, although they may possess limited capacity to adapt to rapidly changing environments and are thus potentially less resilient to effects of anthropogenic-induced climate change.

### A high-quality genomic resource for one of the world’s rarest birds

Obtaining genomic resources for critically endangered species is often constrained by the scarcity of biological material. This is particularly true for *A. insignis*, whose global population is estimated to comprise only a few dozen mature individuals inhabiting remote Himalayan river systems. Our highly contiguous assembly, with a contig N50 exceeding 4.6 Mb, 95.3% complete BUSCO genes and chromosome-scale scaffolding covering 99.5% of the genome, therefore represents an important resource beyond the scope of the present study. It provides a reference for future population genomic investigations, comparative studies within Ardeidae, and conservation genetic monitoring of remaining populations.

The phylogenomic analyses further demonstrate the value of combining mitochondrial and nuclear genomic data. Both datasets consistently place *A. insignis* as the sister species to *A. purpurea*, corroborating previous data based on mitochondrial sequences only (Duan et al. 2018). However, in contrast to Duan et al. (2018), we recovered only the canonical set of 13 mitochondrial protein-coding genes characteristic of avian mitogenomes (Desjardins and Morais 1990; Mindell et al. 1998), suggesting that the additional gene previously reported in Duan et al. (2018) represents an annotation error.

Although the placement of *A. insignis* was fully concordant between the mitochondrial and the nuclear datasets, relationships within the large-heron (*Ardea*) radiation differed slightly. Similar mitonuclear discordance has been previously reported for the Ardeidae family (Hruska et al. 2023) and is increasingly recognized as a common feature of recent avian radiations. Mitonuclear discordance may arise through incomplete lineage sorting or historical introgression among closely related species, or the different inheritance patterns of mitochondrial and nuclear genomes (Edwards et al. 2016). While resolving these evolutionary relationships was not the primary aim of our study, our analyses provide an important genomic foundation for future comparative genomic investigations of Ardeidae evolution.

### A long history of small effective population size rather than recent genomic collapse

Perhaps the most striking result of our study is the remarkable consistency among multiple independent genomic indicators of long-term demographic history. GenomeScope estimated exceptionally low heterozygosity, both *A. insignis* individuals exhibited markedly reduced genome-wide genetic diversity compared with the widespread *A. herodias*, and PSMC analyses consistently reconstructed prolonged periods of reduced effective population size in *A. insignis*. Importantly, these demographic trajectories remained robust across alternative PSMC parameterizations specifically designed to avoid recently identified methodological artefacts (Hilgers et al. 2025), increasing confidence that they reflect genuine evolutionary history rather than analytical bias.

Runs of homozygosity (ROHs) provide additional insight into the temporal scale of these demographic processes. Both *A. insignis* individuals contained substantially longer ROHs than either *A. herodias* individual, consistent with a prolonged history of reduced effective population size. In both *A. insignis* genomes, short ROHs (<1 Mb), which generally reflect older autozygosity accumulated under long-term small population sizes, were abundant (Ceballos et al. 2018). The two individuals nevertheless differed in their burden of longer ROHs (1–5 Mb), which indicate more recent shared ancestry between parental chromosomes: “Wing” carried 115 such tracts spanning 209.3 Mb, compared with 48 tracts spanning 69.7 Mb in “Wire” (Figure 3B, D). Genome-wide heterozygosity recovers the same ranking, being roughly threefold lower in “Wing” than in “Wire”, so that both measures identify “Wing” as the more inbred of the two birds. By contrast, long ROHs were absent from both *A. herodias* genomes, and no ROH exceeding 5 Mb was detected in either *A. insignis* individual. Together, these findings suggest that chronically small effective population sizes are a long-standing characteristic of *A. insignis*, while the extent of recent inbreeding differs between individuals.

Interestingly, these genomic patterns contrast with current field observations. The individual with the highest burden of long ROHs (“Wing”) originated from a region in which relatively many active nests have recently been documented, whereas “Wire” originated from a region supporting a comparatively small breeding population and carried substantially fewer long homozygous segments. Although these observations remain preliminary given the limited sample size, the discordance between genomic patterns and contemporary breeding density indicates that recent local demographic processes are not yet reflected in the genomes of surviving birds.

This discordance is expected on temporal grounds. The length of a homozygous tract is inversely related to the number of generations since the common ancestor from which it was inherited, so the 1–5 Mb tracts observed here reflect inbreeding loops several generations in the past rather than in the present one. Consistent with this, no ROH longer than 5 Mb was detected in either bird, indicating that neither individual descends from a recent mating between close relatives. The transformation of Bhutanese river systems over the past few decades is therefore unlikely to have left a detectable imprint on these genomes yet, and the difference between “Wing” and “Wire” is better read as variation in the ancestry of two individuals than as a signal of contemporary breeding density in their respective basins.

Consequently, the genomic signature observed in *A. insignis* is consistent with a species that has persisted at relatively small effective population sizes over evolutionary timescales. Whether the recent fragmentation of its river habitat has already begun to increase local inbreeding cannot be resolved from two genomes, and the ROH data provide no evidence that it has.

### Long-term persistence despite genomic erosion

Small population size is often equated with inevitable extinction. However, recent advances in conservation genomics have demonstrated that this relationship is considerably more nuanced. Several species have persisted over long evolutionary timescales despite extremely low genetic diversity and extensive homozygosity. These include island species such as Berthelot’s pipit (*Anthus berthelotii)* from the madeiran archipelago (Martin et al. 2023) or the Island Fox (*Urocyon littoralis*), which exhibits extraordinarily low genome-wide diversity yet provides evidence that strongly deleterious recessive mutations have been partially removed through long-term purifying selection (Robinson et al. 2018). Similar conclusions have emerged for the vaquita (*Phocoena sinus*), where genomic analyses suggest that long-term rarity may have reduced the frequency of strongly deleterious alleles prior to the recent anthropogenic population collapse (Morin et al. 2021), and for the North Atlantic Right Whale (*Eubalaena glacialis*), where genomic data indicate substantial historical purging despite persistent small population sizes (Orton et al. 2024). Together, these studies illustrate that prolonged demographic persistence can fundamentally alter the genomic consequences of inbreeding. The genomic history reconstructed here therefore suggests that the *A. insignis* is not simply the product of a recent demographic collapse. Instead, it appears to represent a lineage that has persisted through extended periods of demographic adversity, indicating that long-term rarity alone has not prevented its survival.

### Implications for conservation

Although our findings provide encouraging evidence that *A. insignis* has persisted despite prolonged small population sizes, they should not be interpreted as evidence that genetic processes no longer matter. On the contrary, the exceptionally low heterozygosity observed throughout the genome indicates that the species possesses very limited standing genetic variation. Such reduced diversity is expected to constrain adaptive potential and may limit the capacity of populations to respond to emerging pathogens, environmental disturbances, or the rapid climatic changes currently affecting Himalayan river ecosystems (Allendorf et al. 2010; Kardos et al. 2021).

Consequently, the greatest genetic threat facing *A. insignis* may not be historical inbreeding itself but the interaction between limited evolutionary potential and accelerating environmental change. A species that has remained well adapted to relatively stable ecological conditions for thousands of years may struggle to respond to rapid habitat degradation occurring over only a few decades. Under such circumstances, natural selection may simply operate too slowly to compensate for the pace of anthropogenic change.

These findings reinforce the importance of habitat protection as the primary conservation strategy for *A. insignis*. Preserving and restoring suitable riverine breeding habitats, maintaining connectivity among breeding areas, and preventing further population fragmentation address the proximate drivers of decline and are therefore likely to yield the greatest conservation return. Our results also have direct implications for the ex-situ programme now underway. Because the species’ low diversity appears to be largely ancestral rather than recently acquired, conservation breeding is unlikely to restore variation that has not been present for much of the species’ evolutionary history; its value lies instead in buffering demographic stochasticity and safeguarding the variation that remains. In this context, the difference in long ROH between our two individuals is informative, indicating that recent inbreeding is not uniform among the remaining individuals and that genomic screening of founders and prospective pairs could limit further accumulation of inbreeding in captive and reinforced populations. At the same time, genomic monitoring should become an integral component of future conservation efforts, allowing changes in genetic diversity, inbreeding, and connectivity to be tracked as management actions are implemented.

In conclusion, our study demonstrates that *A. insignis* has survived a long evolutionary history of small population size and genomic erosion. Its fate has therefore not been determined by demographic history alone. Rather, the future persistence of this critically endangered species will likely depend on whether conservation efforts can preserve the ecological conditions under which it has persisted over evolutionary timescales. The challenge facing *A. insignis* may thus be less one of recovering from its evolutionary past than of surviving the unprecedented environmental changes of the Anthropocene. In practical terms, this means safeguarding sufficiently large, connected riverine landscapes and adjacent breeding forests capable of supporting viable populations into the future.

## Authors contributions

Martin Kapun: Conceptualization, Data curation, Formal analysis, Methodology, Project administration, Resources, Software, Validation, Visualization, Writing – Original Draft; Tshering Tobgay: Project administration, Conceptualization, Investigation, Funding acquisition, Resources, Writing – Original Draft; Alexandra Wanka: Data curation, Investigation, Methodology, Resources, Validation, Writing – Review & Editing; Wolfgang Fiedler: Investigation, Funding acquisition, Project administration, Writing – Review & Editing; Tricia C. Goulding: Investigation, Methodology, Writing – Review & Editing; Andreas Kroh: Formal analysis, Methodology, Resources, Validation, Writing – Review & Editing; Luise Kruckenhauser: Resources, Validation, Writing – Review & Editing; Samten Leki: Resources, Investigation, Supervision, Writing – Review & Editing; Thinley Phuntsho: Resources,, Investigation, Writing – Review & Editing; Marcela Suarez-Rubio: Resources, Validation, Funding acquisition, Investigation, Map, Writing – Review & Editing; Sonam Tshering: Resources, Investigation, Writing – Review & Editing; Swen C. Renner: Project administration, Conceptualization, Funding acquisition, Resources, Validation, Investigation, Writing – Original Draft;

## Permits and ethical considerations

All fieldwork was conducted in Bhutan. Capture, handling and blood sampling were carried out under Research Permit No. 875227 (Department of Forests and Park Services). Access to and international transfer of the genetic resources were authorised by the National Biodiversity Centre of Bhutan in accordance with the Nagoya Protocol under permits NBC/BRD/7/2023-2024/4179 (issued 28 May 2025) and NBC/BRD/7/2025-2026/1157 and /1148 (both issued 7 April 2026). Sampling was performed by trained personnel, blood volumes were kept to the minimum required for genomic analysis, and no bird was harmed for the purposes of this study. Both birds were sampled alive. “Wing” remains alive in captivity and is permanently non-releasable after partial amputation of a wing following a collision and severe infection; “Wire” died in July 2025 from a collision with a power line, a death unrelated to this study.

## Acknowledgments

We thank the Royal Society for Protection of Nature for logistical and institutional support throughout this work, and Dr. Kinley Tenzin for his unconditional confidence in the team, which comfortably preceded any evidence that the project would work. We are very grateful to our rafters, who got the field team across the river and, rather more importantly, back again with the sample and the sampling team still aboard and alive.

## Data availability

The raw sequencing data and the assembled genome are publicly available through NCBI under BioProject accession PRJNA1505165. All custom scripts and code used for the bioinformatic analyses are available on GitHub at https://github.com/capoony/ArdeaInsignis_PopGen.

Large language models were used as research assistance during this study. Claude Sonnet v.5 supported code generation, code review, and documentation of bioinformatic workflows, while Claude Opus v5.0 was used for language editing, stylistic revision, and proofreading of the manuscript. All analytical decisions, code, results, and final manuscript content were reviewed and validated by the authors.

