## Supplementary Figure and Methods for "The fate of a dynasty: Population genomics uncovers the demographic history of *Ardea insignis*, one of the rarest bird species in the world"

### Supplementary Materials

#### Supplementary Figures

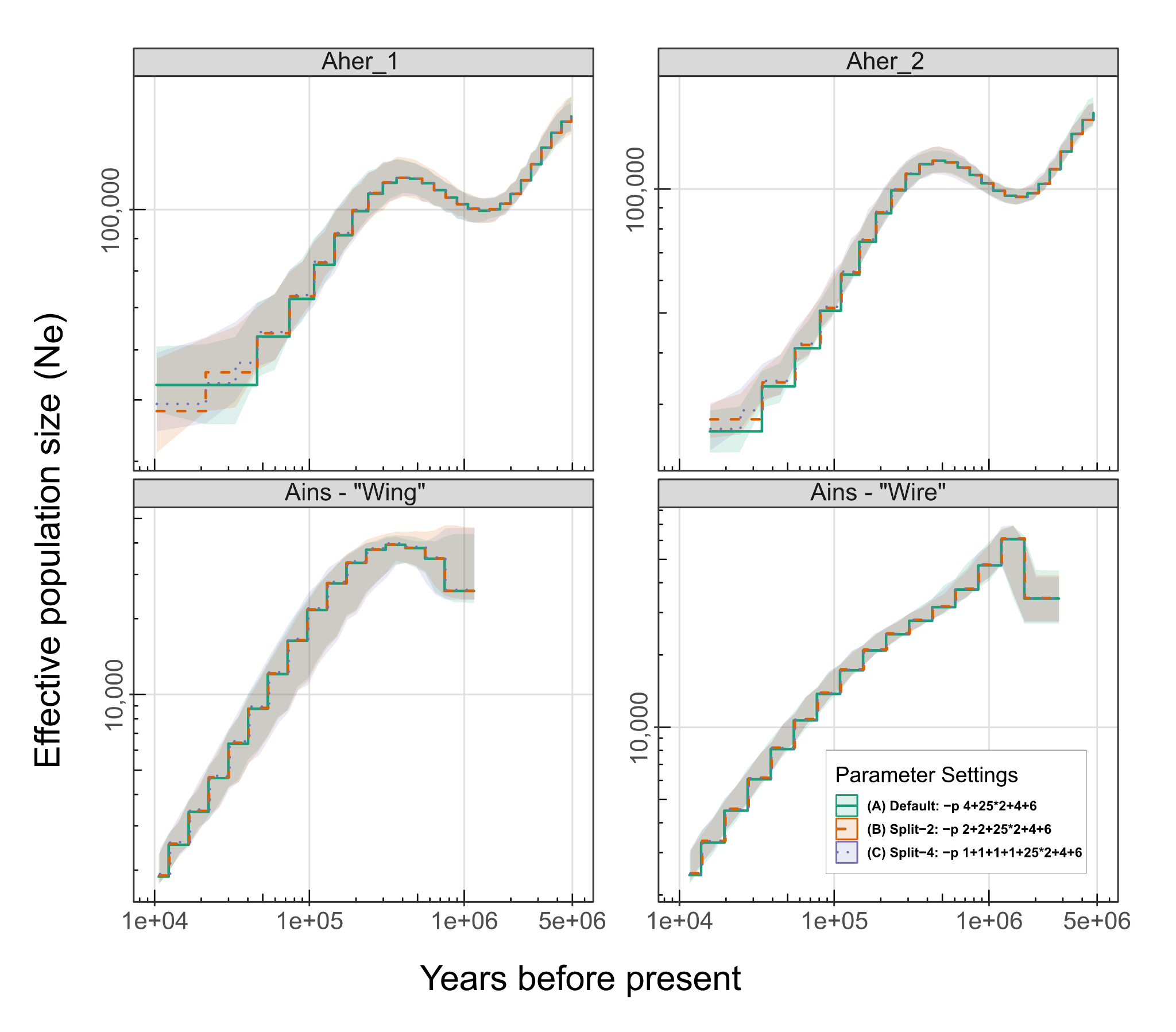

**Figure S1.** Comparison of PSMC demographic reconstructions across alternative time interval parameterizations. Historical effective population size (Nₑ) trajectories inferred with PSMC for two *Ardea insignis* and two *Ardea herodias* individuals using three alternative time interval parameterizations: (A) default (-p 4+25*2+4+6), (B) Split-2 (-p 2+2+25*2+4+6), and (C) Split-4 (-p 1+1+1+1+25*2+4+6). Solid lines represent the maximum-likelihood PSMC estimates, and shaded areas indicate the 95% confidence intervals derived from 100 bootstrap replicates. Demographic trajectories were scaled assuming a generation time of 10 years and a neutral mutation rate of 1.23 × 10⁻⁸ mutations per site per generation. All genomes were mapped to the *A. insignis* reference genome to ensure analytical consistency. The close agreement among parameterizations demonstrates that the inferred demographic histories are robust to the choice of PSMC interval scheme. Consistent with the recommendations of Hilgers et al. (2025), demographic features were considered reliable only when recovered across the modified parameterizations (B and C), whereas patterns unique to the default setting were interpreted cautiously as potential methodological artefacts.

#### Supplementary Methods - Mitochondrial phylogeny

##### Taxon selection

To construct a mitochondrial phylogeny, all currently available Ardaeidae RefSeq mitogenomes were downloaded from GenBank (last checked 31.7.2026). *Ciconia ciconia* (White stork) (Acc.No. NC_002197.1 = AB026818.1, Yamamoto 1999, unpublished) was used as an outgroup.

The RefSeq sequence for *Egretta garzetta* (Acc.No. NC_023981.1 = KJ192197.1, Zou et al. 2015) was found to contain deletions in the ND6 gene (Zou et al. 2014 already mentioned that “its base composition was very different from the other 12 PCGs”) and thus replaced in the analysis by a more complete mitogenome sequence of the same species (OL335994.1, Luo 2021, unpublished)

##### Mitogenome assembly

The mitochondrial genome of *A. insignis* was extracted directly from the assembled genome using MitoFinder v1.4.1 (Allio et al., 2020), employing the mitochondrial genome of the Great Blue Heron (*Ardea herodias*) as the annotation reference. This draft mitogenome, however, could not be circularized (missing bases 11,551 to 11.561 of the published *A. insignis* mitogenome, Acc.No. NC_040004.1, Duan et al. 2018) and showed regions of zero read depth at the end of the control region when mapping the trimmed reads back to the draft mitogenome sequence using Bowtie2 v2.4.4 (Langmead & Salzberg 2012) using the parameters --very-sensitive, --end-to-end, --no-mixed and --no-unal and read counts were called using the samtools v1.12 (Danecek et al. 2021) function depth.

Consequently, trimmed Illumina reads of the *A. insignis* individuals “Wing” and “Wire” were mapped against the published *A. insignis* mitogenome using Bowtie2 v2.4.4 and subsequently assembled *denovo* using SPAdes v3.15.3 (Prjibelski et al. 2020). As can be seen by the aligned contigs, while most of the mitogenome is covered by a single large contig with high read depth, a highly repetitive region with numerous tandem repeats with a length of 22 bp and TAACAAATTAACGAATAACAGR (as checked by the online implementation of TandemRepeatsFinder v4.0.9 (Benson 1999) at https://tandem.bu.edu/trf/home (last checked 1.8.2026). This tandem repeat region is also present in the published *A. insignis* mitogenome where it shows a copy number of 12.7 (the last repeat being truncated to TAACAAATTAACGAA). In addition, individual “Wire” showed three contigs with low read depth that are similar to mitochondrial ND3, COX2-ATP6, and ND4 regions, but highly divergent from the published *A. insignis* mitogenome. Instead, their closest NCBI nucleotide BLAST (Camacho et al. 2009; Priyam et al. 2019) matches are sequences from *Gavia arctica*, *Phoenicopterus roseus*, and *Puffinus yelkouan* respectively. These contigs are here interpreted as NUMTs (Lopez et al. 1994).

Due to the inability of the Illumina data to unambiguously resolve the repetitive region, nanopore reads mapping to the draft mitogenome of “Wing” were identified by read mapping with Bowtie2 v2.4.4. The resulting 786 reads were then assembled with Flye v2.9 (Kolmogorov et al. 2019) using standard the –nano-raw and –nano-hq modes. Both assemblies generated a single contig of 19,342 bp each with identical sequence for both assembly modes. The contig could be circularized and – ignoring copy number variation of the tandem repeat – is 99.89 % identical to the published *A. insignis* mitogenome (Acc.No. NC_040004.1) of Duan et al. (2018). The tandem repeat region at the end of the control region in this assembly shows a copy number of 43.7 (the last repeat again being truncated to TAACAAATTAACGAA). This assembly is here considered the best approximation of the *A. insignis* mitogenome of the individual “Wing”. For “Wire” the copy number of the tandem repeat region could not be unambiguously resolved but is at least 5.7 the last repeat again being truncated to TAACAAATTAACGAA). In addition, in “Wire” only an incomplete mitogenome could be recovered from the generated sequence data missing positions 15,487 to 18,374 in comparison to the published *A. insignis* mitogenome (Acc.No. NC_040004.1), which largely corresponds to the second copy of the duplicated control region. Apart from the missing portion, the mitogenome of “Wire” is almost identical to “Wing” (sequence identity 99.99%) and closely similar to NC_040004 (99.89%).

Analysis of the tandem repeats matching the repeat consensus sequence TAACAAATTAACGAATAACAGR in the nanopore reads fully spanning the repeat region of the mitogenome of “Wing” reveals intra-individual copy number variation between 26 and 91 (mean: 55.3, median: 52, n=173). The custom script employed is available under <https://github.com/a-kroh/MitoTools/blob/main/count_repeats.py>. Such intra-individual tandem repeat copy number variation has also been documented in the mitochondrial genomes of fish (Kijewska et al. 2009), various insects (Chen et al. 2020; Ye et al. 2025) and well documented in the control region of various birds (Berg et al. 1995)

##### Mitogenome annotation

For annotation MITOS2 v2.1.10 (Bernt et al. 2013; Donath et al. 2019) as implemented on the Galaxy Platform (Instance: usegalaxy.eu; The Galaxy Community et al. 2024) using mitfi (Jühling et al. 2012) in glocal and local mode (--locandgloc) was employed. MITOS2 annotations and specifically individual gene boundaries were manually verified and – where necessary – adjusted by aligning the sequences for all individual genes with those of the other Ardeidae species used in the phylogenetic analysis (Table S21), by confirming that PCGs annotations did not produce stop codons and that tRNAs showed the canonical cloverleaf secondary structures.
